# Diverse commensal *E. coli* clones and plasmids disseminate antimicrobial resistance genes in domestic animals and children in a semi-rural community in Ecuador

**DOI:** 10.1101/581967

**Authors:** Liseth Salinas, Paúl Cárdenas, Timothy J. Johnson, Karla Vasco, Jay Graham, Gabriel Trueba

## Abstract

The increased prevalence of antimicrobial resistance (AMR) among Enterobacteriaceae has had major clinical and economic impacts in human medicine. Many of the multi-drug resistant (MDR) Enterobacteriaceae found in humans are community-acquired and linked to food animals (i.e. livestock raised for meat and dairy products). In this study, we examined whether numerically dominant, commensal *Escherichia coli* strains from humans (n=63 isolates) and domestic animals (n=174 isolates) in the same community and with matching phenotypic AMR patterns, were clonally related or shared the same plasmids. We identified 25 multi-drug resistant isolates (i.e. resistant to 3 or more antimicrobial classes) that shared identical phenotypic resistance patterns. We then investigated the diversity of *E. coli* clones, AMR genes and plasmids carrying the AMR genes using conjugation, replicon typing and whole genome sequencing. None of the MDR *E. coli* isolates (from children and domestic animals) analyzed were clonal. While the majority of isolates shared the same antimicrobial resistance genes and replicons, DNA sequencing indicated that these genes and replicons were found on different plasmid structures. Our findings suggest that nonclonal resistance gene dissemination is common in this community and that diverse plasmids carrying AMR genes presents a significant challenge for understanding the movement of AMR in a community.

**IMPORTANCE:** Even though *Escherichia coli* strains may share nearly identical AMR profiles, AMR genes, and overlap in space and time, the diversity of clones and plasmids challenges to research that aims to identify sources of AMR. Horizontal gene transfer appears to play a much larger role than clonal expansion in the spread of AMR in the community.

## INTRODUCTION

Antimicrobial resistance (AMR), especially among Enterobacteriaceae, constitutes an increasing threat to global health (1, 2). Some of the bacterial AMR found in humans has been linked to food-animals (i.e. livestock raised for meat and dairy products) (3). Studies have documented that antimicrobial use in food-animal production is a regular practice in Ecuador and many other countries across the globe (4–7), and that this use increases the likelihood of both the presence of multi-drug resistant (MDR) bacteria in the human microbiota and horizontal gene tranfer of AMR genes to human microbiota (8–10).

Increases in AMR may be greater in low- and middle-income countries (LMICs) than in high-income countries in-part because of the lack of regulatory agencies controlling the use of antimicrobials for humans and food-animals (11, 12). Additionally, contact with food-animal waste, a potential reservoir of drug-resistant bacteria and mobile genenic elements associated with AMR genes (13), can be higher in food-animal producing regions of LMICs than industrialized countries as untreated food-animal wastes are often used to fertilize crops (14). Most research on AMR transmission associated with food animals has focused on commercial-scale production (3, 15, 16), and little research has focused on small-scale food-animal production which is increasingly found to use antimicrobials (15). Despite the potential of small-scale food animals to transmit AMR in a community (17–20), this connection is poorly understood.

Understanding the potential for small-scale food-animal production to spread AMR to human microbiota is critical (21, 22). A study looking at AMR genes in a community suggests that a person’s habitat explains the variation in AMR carriage and that AMR is significantly correlated with the composition of the community and not “randomly distributed across habitats” (19).

*E. coli* is an important species associated with the AMR crisis because it can evolve from a commensal, drug susceptible state to become multi-drug resistant and can cause opportunistic infections (23). Due to its abundance in the intestine, its ability to grow in fecal matter outside the host, and its ability to colonize different hosts, *E. coli* is probably the among most common members of the microbiota transmited beween mammals (24). *E. coli* also is very active in the horizontal transer of AMR genes to other bacteria. The majority of previous studies examining AMR *E. coli* have analyzed colonies isolated in selective media with antimicrobials. In this study, we analyzed numerically dominant *E. coli* clones (isolated in plates without antibiotics), characterized as the clones that were present in the highest proportion, obtained from the fecal samples from children and domestic animals. The goal of this study was to better understand the *E. coli* population dynamics in a community and how AMR can spread within the community (25). Understanding how AMR genes, and its associated clones, can spread in a community between humans, livestock and poultry has the potential to inform policies that aim to mitigate the rise in AMR.

## MATERIALS AND METHODS

### Study location

The study was carried out in the semi-rural community of Otón de Vélez, in the parish of Yaruquí located at an altitude of approximately 2,500 meters above sea level, east of the capital city of Quito. Inhabitants in this community commonly practice small-scale food animal production. Sixty-five households were recruited randomly and were enrolled in the study if they met the inclusion criteria.

Stool samples were obtained from 64 children and 203 fecal samples from 12 species of domestic animals. Of the sampled households, 68% had chickens, 64.5% guinea pigs (raised for food), 64.5% dogs, 58% pigs, 32.3% rabbits, 11.3% cattle and cats, ducks, quails, sheep, geese, horses were present in 10%, 8%, 5%, 3%, 1.6% and 1.6% of the households.

### Ethical considerations

The study protocol was approved by the Institutional Animal Care and Use Committee at the George Washington University (IACUC#A296), as well as the Bioethics Committee at the Universidad San Francisco de Quito (#2014-135M) and the George Washington University Committee on Human Research Institutional Review Board (IRB#101355).

### Sample collection

Fecal samples were collected from children less than five years of age and from domestic livestock, poultry and pets living in the children’s household, from June to August 2014. Stool samples from the children were collected by the child’s primary caretaker (we provided instructions to the caretakers) and animal fecal matter was collected by the study team from the environment where the animal was housed or defecated and avoiding fecal matter with potential contamination. The samples were transported in a cooler (approximately 4 °C) to the laboratory and were processed within eight hours of collection.

### *E. coli* Isolation

Fecal samples were streaked onto MacConkey Lactose agar medium and incubated at 37 °C for 18 hours without antimicrobials, after which at least one lactose positive colony was transferred to Chromocult® Coliform agar for the identification of *Escherichia coli*, through its β-D-glucuronidase activity. *E. coli* isolates were preserved at −80 °C in Brain Heart Infusion (BHI) broth with 20% glycerol.

### Antimicrobial Susceptibility Testing

Each isolate was regrown on Nutrient agar at 37 °C for 18 hours for evaluation of antimicrobial susceptibility by disk diffusion method using Mueller-Hinton agar plates according to the resistance or susceptibility interpretation criteria from Clinical and Laboratory Standards Institute (CLSI) guidelines (26). Antimicrobials used for susceptibility testing included: amoxicillin-clavulanic acid (AMC; 20/10 μg), ampicillin (AM, 10 μg), cefotaxime (CTX, 30 μg), cephalothin (CF, 30 μg), chloramphenicol (C; 30 µg), ciprofloxacin (CIP; 5 μg), gentamicin (CN; 10 μg), imipenem (IPM; 10 μg), streptomycin (S; 10 μg), sulfisoxazole (G; 250 μg), tetracycline (TE; 30 µg) and trimethoprim-sulfamethoxazole (SXT; 1.25 / 23.75 μg) (26).

### Bacterial Conjugation

Following antimicrobial susceptibility testing, we selected 25 MDR isolates that shared antibiotic resistance to 3 antimicrobials: TE-G-SXT, although some isolates showed additional antimicrobial resistance (Table 2); this was the most common multiresistance pattern in the community. Isolates were selected based on identifying similar MDR phenotypic patterns present in isolates from domestic animals only, humans only, and both sources. Each one of the 25 MDR isolates were used as donor strains for the conjugation assay, with three different strains, *E. coli* J53, resistant to sodium azide, *E. coli* TOP10 (Invitrogen, Carlsbad, USA) resistant to rifampicin and *E. coli* TOP10 resistant to nalidixic acid, as receptors. Selection of mutant *E. coli* TOP10, resistant to rifampicin and nalidixic acid, as previously described (27). Prior to each experiment for conjugation, the donor and recipient strains were inoculated in 10 mL of Trypticase Soy Broth (TSB) and grown at 37 ºC for 18 hours, the strains in a logarithmic phase were mixed and incubated at 37 ºC for 18 hours. For the selection of transconjugants, 100 μL of the mix was inoculated by spread plate method onto Nutrient agar supplemented with tetracycline (15 μg/mL) and one of the following antimicrobials: sodium azide (200 μg/mL) (28), rifampicin (100 μg/mL) or nalidixic acid (30 μg/mL) according to the recipient strain used (27). Tetracycline was used as donor selector, since this resistance was present in all of 25 MDR isolates. The transconjugants were evaluated for the 12 antimicrobials tested by the disk diffusion method in order to determine the acquired resistance. The transconjugants were stored as described above.

### DNA Extraction

Total DNA was extracted from the same 25 isolates showing similar AMR phenotypic profiles using the DNeasy® Blood & Tissue Kit (Qiagen, Hilden, Germany) following the manufacturer's recommendations. Plasmid DNA from the transconjugants was extracted using QIAprep Spin Miniprep Kit (Qiagen, Hilden, Germany) following the manufacturer's recommendations.

### Replicon Typing

Replicon typing of transconjugants was carried out from plasmid DNA, using PCR-based replicon typing kit (PBRT Kit; Diatheva, Cartoceto, Italy) following the manufacturer’s instructions (29).

### DNA Sequencing

For the 25 isolates selected, whole genome sequencing (WGS) was performed using Illumina MiSeq. Sequencing was performed at the University of Minnesota Mid-Central Research and Outreach Center (Willmar, MN) using a single 250-bp dual-index run on an Illumina MiSeq with Nextera XT libraries to generate approximately 30- to 50-fold coverage per genome. Several isolates were also sequenced using PacBio technology at the University of Minnesota Genomics Center (Minneapolis, MN). SMRTbell template libraries were generated from previously isolated unsheared raw genomic DNA using Pacific Biosciences SMRTbell template preparation kit 1.0 (Pacific Biosciences, Menlo Park, CA). Finished DNA libraries subsequently were subjected to DNA size selection using the BluePippin DNA size selection system (Sage Science Inc.) with a 7-kb cutoff to select DNA fragments greater than 7 kb. Sequencing was performed on the PacBio Sequel (Pacific Biosciences, Menlo Park, California).

### Data Analysis

Illumina raw reads were quality-trimmed and adapter-trimmed using trimmomatic (30). Reads were assembed using SPaDES assembler (31). PacBio reads were assembled using canu (32). Contigs obtained were then annotated with Prokka (33). Resistance genes (ResFinder 2.1), Gram negative plasmid types (PlasmidFinder 1.3), plasmid allele types (pMLST; pMLST 1.4), and multilocus sequence typing profiles, (MLST; MLST 1.8) were obtained based on the sequences obtained using the Center for Genomic Epidemiology tools (http://www.genomicepidemiology.org/) (34). Phylogenetic analysis of individual genes/segments was performed using MEGA7 (35), whilst WGS aligments were performed using Mauve (36, 37). To visualize the STs and the micro-evolutionary processes, a minimum spanning tree was constructed using PHYLOViZ 2.0 (38). Insertions sequence elements was performed using ISFinder (39). Plasmid maps were constructed using XPlasMap version 0.96 (http://www.iayork.com/XPlasMap/).

## RESULTS

Two-hundred and thirty-seven *E. coli* isolates were recovered, 63 from child fecal samples and 174 from domestic animals. More than one-third of the isolates (38.4 %) were susceptible to all twelve antimicrobials evaluated; 46 (19.4 %) were resistant to one antimicrobial; 19 (8.0%) to 2 antimicrobials, and 81 isolates (34.2%) were multidrug resistant (i.e. resistant to three or more antimicrobial classes).

### Antimicrobial Resistance in Humans and Domestic Animals

In general, isolates obtained from children showed higher phenotypic resistance compared with isolates from domestic animals (Table 1), including resistance to tetracycline (50.8 %), sulfisoxazole (49.2 %) and ampicillin (49.2 %). The highest percentages of phenotypic resistance in isolates obtained from animals were to tetracycline (39.7%), sulfisoxazole (24.1%) and cephalothin (23%), a first-generation cephalosporin. The most frequent MDR profile, found in 13.6% of the isolates, was tetracycline, sulfisoxazole, ampicillin, streptomycin and trimethoprim-sulfamethoxazole; the majority of these isolates belonged to humans (63.6%) (Table 2).

**TABLE 1.** Prevalence of phenotypically resistant *E. coli* found in fecal samples from children and animals.

| Antimicrobial | Children<br>(n=63) | Domestic Animals<br>(n=174) |
| --- | --- | --- |
| Amoxicillin-clavulanic acid | 6 (9.5%) | 4 (2.3%) |
| Ampicillin | 31 (49.2%) | 35 (20.1%) |
| Cefotaxime | 4 (6.4%) | 10 (5.8%) |
| Cephalothin | 20 (31.8%) | 40 (23.0%) |
| Chloramphenicol | 6 (9.5%) | 32 (18.4%) |
| Ciprofloxacin | 4 (6.4%) | 16 (9.2%) |
| Gentamicin | 2 (3.2%) | 2 (1.2%) |
| Imipenem | 0 (0%) | 2 (1.2%) |
| Streptomycin | 26 (41.3%) | 31 (17.8%) |
| Sulfisoxazole | 31 (49.2%) | 42 (24.1%) |
| Tetracycline | 32 (50.8%) | 69 (39.7%) |
| Trimethoprim-sulfamethoxazole | 27 (42.9%) | 36 (20.7%) |

**TABLE 2.**
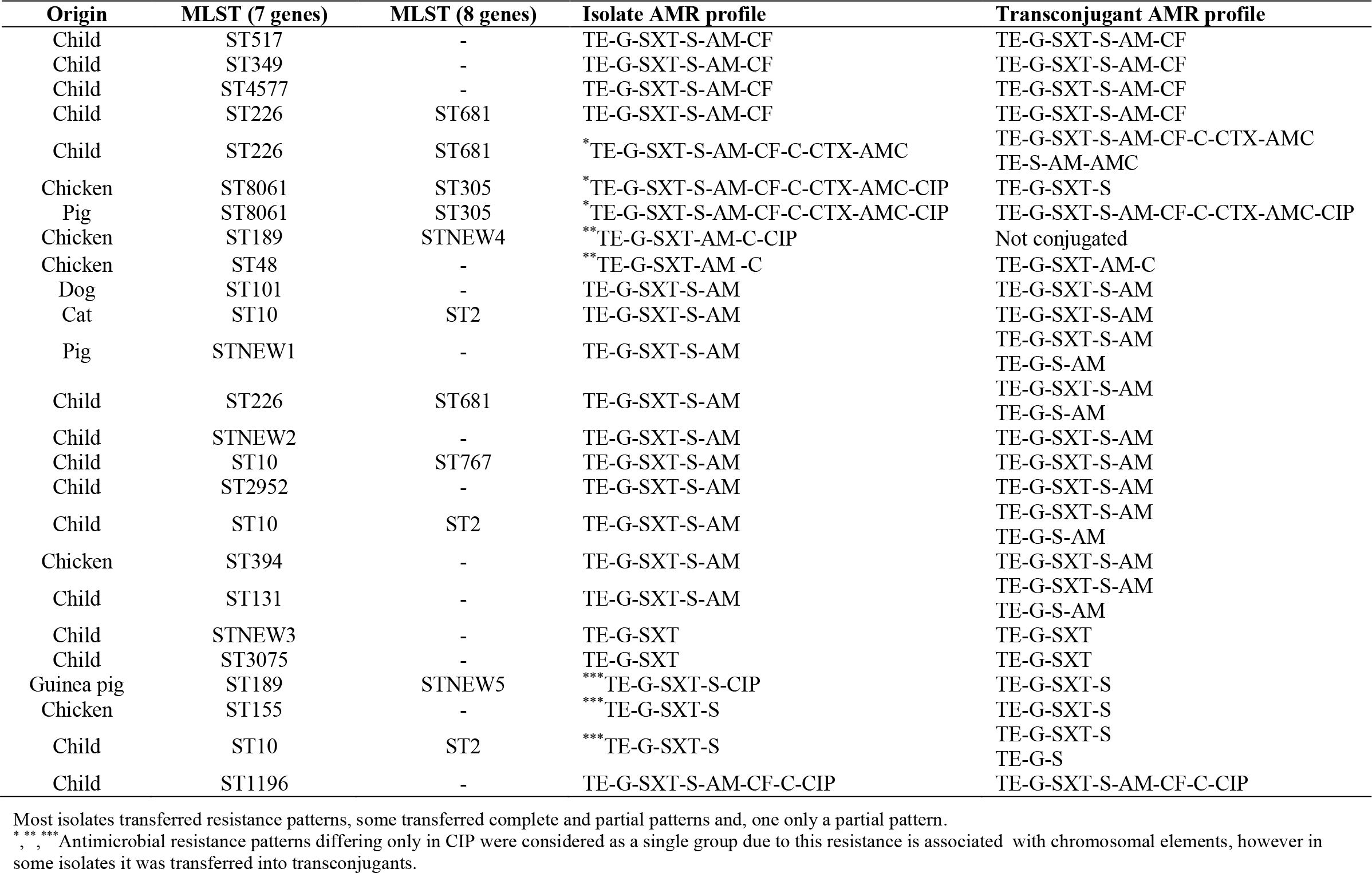
Phenotypic antimicrobial resistance profile of multi-drug resistant *E. coli* isolates (from children and domestic animals) and transconjugants.

### Bacterial Conjugation

For the bacterial conjugation assays, we selected 15 MDR isolates from children and 10 MDR isolates from domestic animals; all of the isolates grouped into same 7 MDR phenotypic patterns as described above. We obtained a total of 30 transconjugants from 24 isolates. Twenty-three isolates transferred their complete phenotypic resistance pattern to the receptor bacteria, and 6 isolates transferred both partial resistance and total resistance and 1 isolate transferred partial resistance only (Table 2).

### Isolate Genotyping

For the 25 selected *E. coli* isolates, we conducted MLST, which sequences internal fragments of the following genes: *adk*, *fumC*, *gyrB*, *icd*, *mdh*, *purA* and *recA.* We identified 18 different STs; seven STs were found in domestic animals only (ST8061, ST189, ST48, ST101, STnew-1 (not available in database), ST394 and ST155), and 10 STs were found in isolates from children only (ST157, ST349, ST4577, ST226, STnew-2 (not available in database), ST2952, ST131, STnew-3 (not available in database), ST3075 and ST1196), and ST10 was present in isolates from both sources (Table 2).

Among animal isolates, an isolate from a guinea pig and another from a chicken, belonged to ST189 and an isolate from chicken and another from pig belonged to ST8061. However, when we conducted extended MLST, which includes additional genes (*dinB, icdA, pabB, polB, putP, trpA, trpB* and *uidA*), the results showed that the isolates were different; isolates identified as ST189 were found to be different: STnew-4 and STnew-5. Upon further analysis, WGS showed that none of the isolates were clonal. Among the human isolates, 3 belonged to ST226, which were similar to ST681. Three human isolates and one from a cat belonged to ST10, however whole genome sequencing showed that the two human isolates shared a more recent common ancestor but were not identical (both human isolates differed by 5,932 SNPs and the human isolates and the cat isolates differed by 6,138-7,409 SNPs) Additional extended MLST analysis of the of ST10 isolates showed that three belonged to ST2 and one to ST767 (Table 2).

### Plasmid Genotyping

We identified 17 replicon types in the transconjugants; 7 replicon types (X3, FIC, I1**γ**, W, X2, B/O, and K) originated in human isolates, 9 (L, P, FIIS, FII, FIA, A/C, **γ**, I2, and FIB) were present in isolates from both human and domestic animals; and, I1**α** was present in only one isolate from an animal. The most common replicons were FII and FIIS, which were found in 23 of the isolates (92%) and 22 (88%) respectively (Table 4).

**TABLE 3.** Antimicrobial resistance genes and antimicrobial resistance profile of *E. coli* islotes (from children and domestic animals)

| Origin | AMR genes | Phenotypic AMR profile |
| --- | --- | --- |
| Child | *bla*<sub>TEM-1B</sub>, *dfrA8*, *qnrB19*, *strA*, *strB*, *tetB* | TE-G-SXT-S-AM-CF |
| Child | *bla*<sub>TEM-1B</sub>, *dfrA7*, *dfrA8*, *strA*, *strB*, *sul1*, *sul2*, *tetA* | TE-G-SXT-S-AM-CF |
| Child | *bla*<sub>TEM-1B</sub>, *dfrA8*, *strA*, *strB*, *sul2*, *tetB* | TE-G-SXT-S-AM-CF |
| Child | *aadA1*, *bla*<sub>TEM-1B</sub>, *dfrA15*, *qnrB19*, *tetA* | TE-G-SXT-S-AM-CF |
| Child | *aadA1*, *bla*<sub>TEM-1B</sub>, *dfrA15*, *qnrB19*, *sul1*, *tetA* | TE-G-SXT-S-AM-CF-C-CTX-AMC |
| Chicken | *aadA1*, *bla*<sub>CMY-2</sub>, *bla*<sub>TEM-1B</sub>, *catA1*, *cmlA1*, *floR*, *fosA*, *lnuF*, *qnrB19*, *strA*, *strB*, *sul2*, *sul3*, *tetA* | TE-G-SXT-S-AM-CF-C-CTX-AMC-CIP |
| Pig | *bla*<sub>CMY-2</sub>, *bla*<sub>TEM-1B</sub>, *catA1*, *qnrB19*, *strA*, *strB*, *sul2*, *tetA* | TE-G-SXT-S-AM-CF-C-CTX-AMC-CIP |
| Chicken | *dfrA14*, *strA*, *strB*, *sul2* | TE-G-SXT-AM-C-CIP |
| Chicken | *aadA1*, *aadA2*, *bla*<sub>TEM-1B</sub>, *cmlA1*, *dfrA12*, *dfrA14*, *mef*, *qnrB19*, *strA*, *strB*, *sul2*, *sul3*, *tetA*, *tetB* | TE-G-SXT-AM-C |
| Dog | *aadA2*, *bla*<sub>TEM-1B</sub>, *dfrA12*, *sul3*, *tetA*, *tetM* | TE-G-SXT-S-AM |
| Cat | *bla*<sub>TEM-1B</sub>, *dfrA8*, *strA*, *strB*, *sul2*, *tetB* | TE-G-SXT-S-AM |
| Pig | *bla*<sub>TEM-1B</sub>, *dfrA8*, *strA*, *strB*, *sul2*, *tetB* | TE-G-SXT-S-AM |
| Child | *bla*<sub>TEM-1B</sub>, *dfrA8*, *strA*, *strB*, *sul2*, *tetA* | TE-G-SXT-S-AM |
| Child | *bla*<sub>TEM-1B</sub>, *dfrA8*, *strA*, *strB*, *sul2*, *tetB* | TE-G-SXT-S-AM |
| Child | *aadA5*, *bla*<sub>TEM-1B</sub>, *dfrA17*, *mphA*, *strA*, *strB*, *sul1*, *sul2*, *tetA* | TE-G-SXT-S-AM |
| Child | *bla*<sub>TEM-1B</sub>, *dfrA8*, *strA*, *strB*, *sul2*, *tetA* | TE-G-SXT-S-AM |
| Child | *bla*<sub>TEM-1B</sub>, *dfrA5*, *strA*, *strB*, *sul1*, *sul2*, *tetB* | TE-G-SXT-S-AM |
| Chicken | *aadA1*, *bla*<sub>TEM-1B</sub>, *dfrA1*, *strA*, *strB*, *sul1*, *sul2*, *tetA* | TE-G-SXT-S-AM |
| Child | *bla*<sub>TEM-1B</sub>, *dfrA8*, *strA*, *strB*, *sul2*, *tetB* | TE-G-SXT-S-AM |
| Child | *aadA5*, *dfrA17*, *qnrB19*, *sul2*, *tetA* | TE-G-SXT |
| Child | *dfrA14*, *strA*, *strB*, *sul2*, *tetA* | TE-G-SXT |
| Guinea pig | *aadA24*, *dfrA14*, *dfrA15*, *strA*, *strB*, *sul1*, *sul2* | TE-G-SXT-S-CIP |
| Chicken | *dfrA14*, *qnrB19*, *strA*, *strB*, *sul2*, *tetA* | TE-G-SXT-S |
| Child | *dfrA5*, *qnrB19*, *strA*, *strB*, *sul1*, *sul2*, *tetA* | TE-G-SXT-S |
| Child | *aadA1*, *aadA2*, *bla*<sub>TEM-1B</sub>, *cmlA1*, *dfrA12*, *floR*, *fosA*, *lnuF*, *sul3*, *tetA* | TE-G-SXT-S-AM-CF-C-CIP |

**TABLE 4.** Plasmid genotyping and antimicrobial resistance profile of isolates (from children and domestic animals) and transconjugants

| Origin | Plasmids | pMLST | Replicons | Transconjugant AMR profile | Isolate AMR profile |
| --- | --- | --- | --- | --- | --- |
| Child | IncFII(pRSB107), IncFIB(AP001918) | FII43, FIB11 | L, P, X3, FIIS, FIC, FII | TE-G-SXT-S-AM-CF | TE-G-SXT-S-AM-CF |
| Child | IncFII(pHN7A8), IncFII, IncQ1 | FI33, FIB29 | P, I1Y, FIIS, FIC, FII | TE-G-SXT-S-AM-CF | TE-G-SXT-S-AM-CF |
| Child | IncFIB(pLF82), IncFII(pHN7A8) | FII11 | FIA, W, A/C, FIIS, X2, FII | TE-G-SXT-S-AM-CF | TE-G-SXT-S-AM-CF |
| Child | IncFII(pSE11), IncFIB(AP001918), IncFII, Col(MG828) | FI79, FIB28 | P, A/C, FIIS, X2, FII | TE-G-SXT-S-AM-CF | TE-G-SXT-S-AM-CF |
| Child | IncFII(pSE11), IncFIB(AP001918), IncFII, Col(MG828) | FI79, FIB28 | P, A/C, FIIS, X2, FII | TE-G-SXT-S-AM-CF-C-CTX-AMC | TE-G-SXT-S-AM-CF-C-CTX-AMC |
| Child | IncFII(pSE11), IncFIB(AP001918), IncFII, Col(MG828) | FI79, FIB28 | A/C, FIIS, X2, FII | TE-S-AM-AMC | TE-G-SXT-S-AM-CF-C-CTX-AMC |
| Chicken | IncB/O/K/Z, IncFII, IncFIB(AP001918), IncQ1 | FII64, FIB27 | FIB, L, P, FIIS, FII | *Not conjugated | TE-G-SXT-S-AM-CF-C-CTX-AMC-CIP |
| Chicken | IncB/O/K/Z, IncFII, IncFIB(AP001918), IncQ1 | FII64, FIB27 | FIB, FIIS, FII | TE-G-SXT-S | TE-G-SXT-S-AM-CF-C-CTX-AMC-CIP |
| Pig | IncFIB(AP001918), IncFII, IncB/O/K/Z, IncQ1 | FII64, FIB27 | FIB, L, P, A/C, FIIS, FII | TE-G-SXT-S-AM-CF-C-CTX-AMC-CIP | TE-G-SXT-S-AM-CF-C-CTX-AMC-CIP |
| Chicken | IncI2, IncY | FI43 | I2, L, A/C, FIIS, FII | **No conjugated | TE-G-SXT-AM-C-CIP |
| Chicken | IncFII(29), IncFIB(K), IncFIA(HI1) | FII29, FIA13 | L | TE-G-SXT-AM-C | TE-G-SXT-AM-C |
| Dog | IncFII, IncN3 | FII34 | L, FIIS, Y, FII | TE-G-SXT-S-AM | TE-G-SXT-S-AM |
| Cat | IncFIB(AP001918), IncFII(pRSB107) | FII1, FIB54 | L, FIIS, Y, FII | TE-G-SXT-S-AM | TE-G-SXT-S-AM |
| Pig | IncFII(pRSB107), IncFIB(AP001918), IncI1 | FII48, FIB 25 | L, FIIS, Y, FII | TE-G-SXT-S-AM | TE-G-SXT-S-AM |
| Pig | IncFII(pRSB107), IncFIB(AP001918), IncI1 | FII48, FIB 25 | I2, FIB, L, P, FIIS, Y, FII | TE-G-S-AM | TE-G-SXT-S-AM |
| Child | IncFIB(AP001918), IncFII(pRSB107), IncFII(29) | FII1, FIB54 | I2, FIB, P, FIIS, Y, FII | TE-G-SXT-S-AM | TE-G-SXT-S-AM |
| Child | IncFIB(AP001918), IncFII(pRSB107), IncFII(29) | FII1, FIB54 | I2, FIIS, FII | TE-G-S-AM | TE-G-SXT-S-AM |
| Child | IncFIB(AP001918), IncFII, IncFII(pRSB107) | FII1, FIB54 | FIIS, FII | TE-G-SXT-S-AM | TE-G-SXT-S-AM |
| Child | IncFIB(AP001918), IncFIA, IncFII(pCoo) | FII10, FIA2, FIB20 | FIB, FIA, A/C, FIIS, FII | TE-G-SXT-S-AM | TE-G-SXT-S-AM |
| Child | IncQ1, IncFII(pCoo) | FII16 | FIIS, FII | TE-G-SXT-S-AM | TE-G-SXT-S-AM |
| Child | IncFII(pRSB107), IncB/O/K/Z, IncI2 | FII6 | FIIS, FII | TE-G-SXT-S-AM | TE-G-SXT-S-AM |
| Child | IncFII(pRSB107), IncB/O/K/Z, IncI2 | FII6 | I2, L, FIIS, FIC, FII | TE-G-S-AM | TE-G-SXT-S-AM |
| Chicken | IncFII(pHN7A8), IncFII, Col(BS512) | FII11 | I1α, FIB, FIA, P, FIIS, FII | TE-G-SXT-S-AM | TE-G-SXT-S-AM |
| Child | IncFIB(AP001918), IncFII, IncFIC(FII), IncFIA, IncFII(pRSB107), IncI1, Col(BS512) | FII1, FIA1, FIB1 | I2, FIIS, FIC, FII | TE-G-SXT-S-AM | TE-G-SXT-S-AM |
| Child | IncFIB(AP001918), IncFII, IncFIC(FII), IncFIA, IncFII(pRSB107), IncI1, Col(BS512) | FII1, FIA1, FIB1 | FIIS, FIC, FII | TE-G-S-AM | TE-G-SXT-S-AM |
| Child | p0111 | NEW1 | I1Y, FIIS, FII | TE-G-SXT | TE-G-SXT |
| Child | IncFII(pCoo), IncY, IncB/O/K/Z | FII43, FIB24 | B/O, I1Y, A/C, FIIS, FIC, FII | TE-G-SXT | TE-G-SXT |
| Guinea pig | IncFIB(AP001918), IncI1, IncI2, Col156 | FII17 | I1α, FIB, P, FII | TE-G-SXT-S | TE-G-SXT-S-CIP |
| Chicken | IncFII(29) | FII29 | FIIS, FII | TE-G-SXT-S | TE-G-SXT-S |
| Child | IncB/O/K/Z, Col(MG828) | NEW2 | P, K | TE-G-SXT-S | TE-G-SXT-S |
| Child | IncB/O/K/Z, Col(MG828) | NEW2 | P, K | TE-G-S | TE-G-SXT-S |
| Child | IncFIB(AP001918), IncFII(29), p0111 | FII29, FIB1 | FIB, FIIS, FII | TE-G-SXT-S-AM-CF-C-CIP | TE-G-SXT-S-AM-CF-C-CIP |

Plasmids and pMLST were obtained from WGS, replicons were characterized from transconjugates. \*Result of replicon typing referred to donor strain, because we obtained only a partial transconjugant. \*\*Result of replicon typing referred to donor strain, because there were no transconjugant obtained.

Whole genome sequencing of the 25 selected isolates identified 22 replicons, and 11 replicons were found in isolates from both children and domestic animals: IncFII(pRSB107), Incl1, IncQ1, IncFII(29), IncY, IncFII, IncB/O/K/Z, Incl2, IncFIB(AP001918), IncFII(pHN7A8) and Col(BS512). Four replicons were found only in animal isolates (IncN3, IncFIA(HI1), IncFIB(K) and Col156), and 7 replicons were found only in human isolates (IncFII(pSE11), IncFII(pCoo), IncFIC(FII), IncFIB(pLF82), Col(MG828), IncFIA, and p0111) (Figure 1a and Table 4).

**FIG 1.**
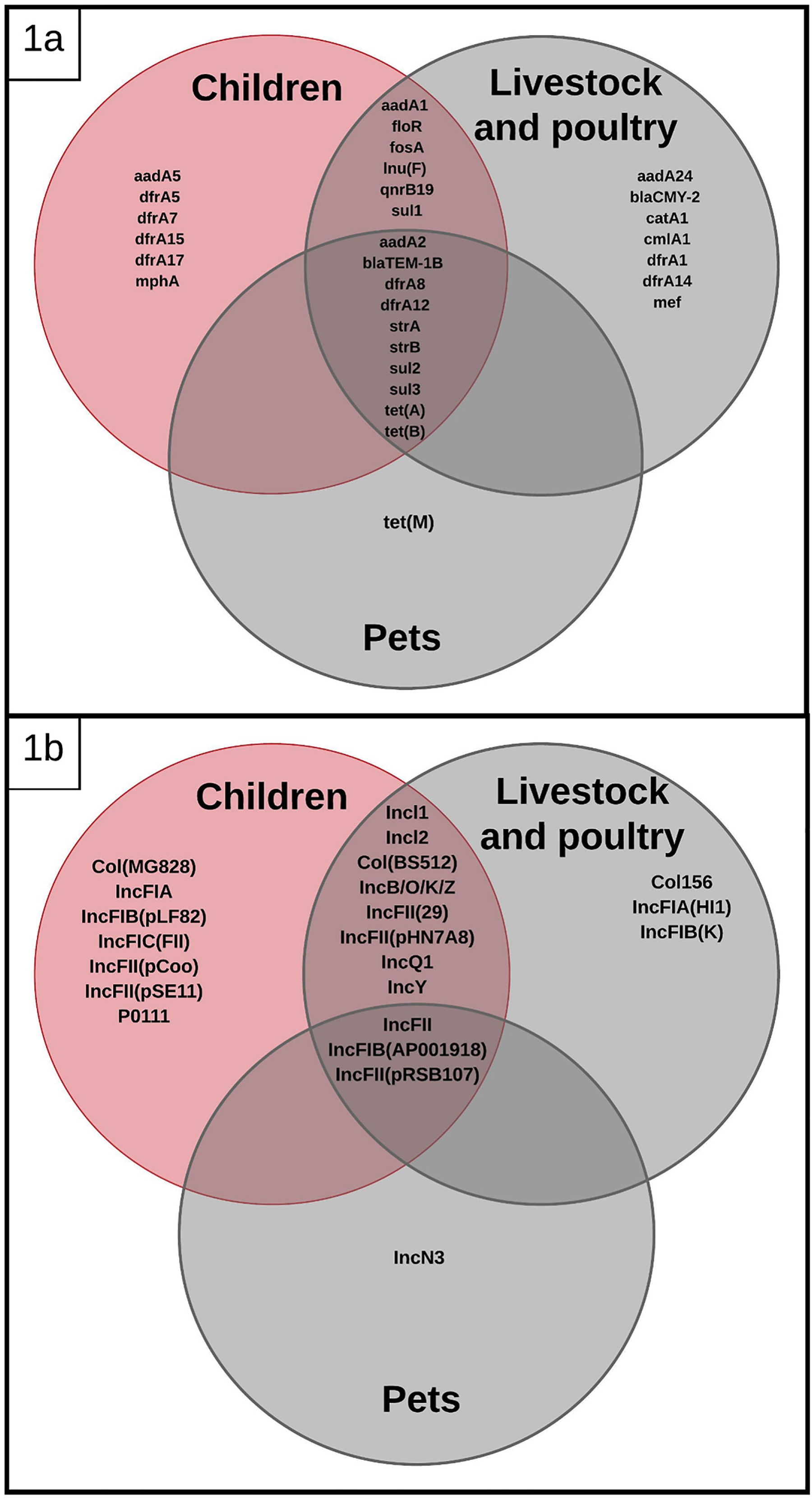
Venn diagrams showing shared antimicrobial resistance genes (1a) and replicons (1b) among the *E. coli* isolates from children, livestock, poultry and pets.

Twenty-eight F-type plasmids were further characterized by pMLST; 8 of them (FII64, FIB27, FI43, FIA13, FII34, FII48, FIB25 and FIB24) came from animal samples, 15 (FIB11, FI33, FI79, FIB28, FII10, FIA2, FIB20, FII16, FII6, FIA1, FII17, FIB1, FIB29, NEW1 and NEW2) from human samples, while 5 (FII43, FII11, FII29, FII1 and FIB54) were identified in both sources. The replicon type did not appear to correlate with a specific phenotypic resistance pattern (Table 4). Further characterization of plasmids using long-read PacBio sequencing demonstrated that none of the plasmids compared were identical based on pMLST or whole plasmid sequence alignments, however there were similarities in AMR genes. For example, the association of Tn*2*(*bla*_TEM-1B_) (with an identical DNA sequence) was encountered in 4 different plasmids from 4 different strains from children in diferent hoseholds. The DNA sequences of Tn*2*(*bla*_TEM-1B_) were identical to others found in other bacterial species in the genbank, suggesting that this association is not novel. Additionally, two different strains (different STs) from two children in different households, had plasmids sharing 73% of gene content but different on replicon types (Supplemental Materials Figure 1).

### Analysis of Antimicrobial Resistance Genes

Whole genome sequencing of the 25 selected isolates showed 30 allelic variants of antimicrobial resistance genes. The following AMR genes were found in children and domestic animals: *bla*_TEM-1B_, *dfr*A8, *dfr*A12, *dfr*A14, *dfr*A15, *qnr*B19, *str*A, *str*B, *tet*A, *tet*B, *sul*1, *sul*2, *sul*3, *flo*R, *aad*A1, *aad*A2, *cml*A1, *Inu*F and *fos*A (Figure 1b). AMR genes found only in domestic animal isolates included: *bla*_CMY-2_, *cat*A1, *mef*, *tet*M, *dfr*A1 and *aad*A24, and AMR genes found only in children included: *dfr*A5, *dfr*A7, *dfr*A17, *aad*A5 and *mph*A (Table 3). Phylogenetic analysis of the most common genes showed that *tet*A*, tet*B and *dfr*A8 were identical; and we found sequences classified as *aad*A1-like*, str*A-like*, str*B-like and *sul*2-like with SNPs, clustering independently from strain origin.

## DISCUSSION

In this semi-rural community, we found that numerically dominant commensal *E. coli* (showing similar antimicrobial resistance and same antibiotic resistance genes) colonizing children and domestic animals in the same period of time and in the same community, are genotypically diverse. We also found that plasmids carrying the same antibiotic resistance genes were distinct. Our research suggests that the a common pool of AMR genes could be co-circulating on different plasmids amongst different *E. coli* clones in a community (Table 3). Even when same resistance gene alleles and same plasmid replicon types were identified across isolates, the plasmids harboring these traits were still distinct. We also found potential evidence of Tn*2* participation in mobility of the gene *bla* _TEM-1B_, as we found this transposon-gene association in 4 different plasmids and 4 genetically different *E. coli*; This is indicative of common pools of transposable elements actively moving genes among different plasmids (51). The findings of this study should raise caution of the conclusions reached in many studies, which have used MLST and replicon typing to identify the sources of AMR genes and have concluded that matching MLST profile and AMR gene profile suggests clonality. Our results are concordant with a recent report showing that WGS of food animal isolates and human bloodstream infections (in the United Kingdom) were different and did not share plasmids, although isolates (from humans and domestic animals) shared some antibiotic resistance genes (40).

Previous studies have found that some numerically dominant *E. coli* strains from domestic animals and humans can be shared within the same household (41–43), although we did not find direct evidence of clone sharing within households enrolled in this study. The reason for this discrepancy may be the reliance, in previous studies, on MLST to detect clonal transmission (44–46), which seems inadequate based on our data and previous reports (47, 48). Another reason may be that we selected individuals in a community instead of individuals in the same household.

We showed evidence that human and domestic animal strains share the same replicons and pMLST profiles: IncFII, IncFII(29), IncFII(pRSB107), IncFII(pHN7A8), IncFIB(AP001918), Incl1, Incl2, IncQ1, IncY, IncB/O/K/Z, Col(BS512), FII1, FII11, FII29, FII43, FIB54. This would suggest that plasmids were shared among numerically dominant and antimicrobial resistant *E. coli* from humans and domestic animals in this community. However, the long read sequencing of plasmids indicated that these plasmids were not identical. Still, allelic variants of some antimicrobial resistance genes were identical among isolates from humans and domestic animals. This again suggests that mobile genetic elements within these diverse plasmids, such as transposons, conjugative transposons, and integrons, may be more actively involved in the mobility of AMR genes between plamids and bacterial cells than plasmid transfer itself (49–51). Also our data suggest that many of the plasmids circulating in *E. coli* (in the same human community) could share many genes (including AMR genes and replicons) but they are not the same.

This remarkable genetic plasticity has been described in some plasmids carrying MDR genes (51, 52). Our report suggest that assessing the real impact dimension of antibiotic use in food animals in public is a very complex endeavor which will be accomplished only through the use of powerful DNA sequencing technology.

## ACKNOWLEDGMENTS

We greatly appreciate the assistance of Valeria Garzon, the Yaruquí community, as well as our colleagues in the Microbiology Institute at the Universidad San Francisco de Quito, in conducting this research.

## Disclaimer

The authors declare no conflict of interest.

**SUPPLEMENTAL MATERIALS FIGURE 1.** Plasmid 6 from sample ECU06 and and plasmid 4 from sample ECU07 alignments on Mauve and gene annotations on Prokka and ISFinder showing shared genomic structure between two different plasmids (different replicon types). Shared genes are colored in green and red, replicon names and locations are in blue boxes, and transposons/insertion sequences are in clear boxes.

## REFERENCES

1. Finley RL, Collignon P, Larsson DGJ, McEwen SA, Li X-Z, Gaze WH, Reid-Smith R, Timinouni M, Graham DW, Topp E. 2013. The scourge of antibiotic resistance: the important role of the environment. Clin Infect Dis 57: 704–710.

2. Savard P, Perl TM. 2012. A call for action: managing the emergence of multidrug-resistant Enterobacteriaceae in the acute care settings. Curr Opin Infect Dis 25: 371–377.

3. Seiffert SN, Hilty M, Perreten V, Endimiani A. 2013. Extended-spectrum cephalosporin-resistant Gram-negative organisms in livestock: an emerging problem for human health? Drug Resistance Updates. 16 SRC-:22–45.

4. Hu Y, Cheng H. 2016. Health risk from veterinary antimicrobial use in China’s food animal production and its reduction. Environ Pollut 219: 993–997

5. Graham JP, Boland JJ, Silbergeld E. 2007. Growth promoting antibiotics in food animal production: an economic analysis. Public Health Rep 79–87.

6. Van Boeckel TP, Brower C, Gilbert M, Grenfell BT, Levin SA, Robinson TP, Teillant A, Laxminarayan R. 2015. Global trends in antimicrobial use in food animals. Proc Natl Acad Sci U S A 112: 5649–5654.

7. Scientific AG on A of the C for MP for VU. 2009. Reflection paper on the use of third and fourth generation cephalosporins in food producing animals in the European Union: development of resistance and impact on human and animal health. J Vet Pharmacol Ther 32: 515–533.

8. Vinueza-Burgos C, Cevallos M, Ron-Garrido L, Bertrand S, Zutter L. 2016. Prevalence and diversity of *Salmonella* serotypes in Ecuadorian broilers at slaughter age. PloS one e0159567

9. Mesa RJ, Blanc V, Blanch AR, Lavilla S, Miro E, Muniesa M, Saco M. 2006. Cortés P, González JJ, Tórtola MT. Extended-spectrum beta-lactamase-producing Enterobacteriaceae in different environments (humans, food, animal farms and sewage). J Antimicrob Chemother 58 SRC-:211–215.

10. Marshall BM, Levy SB. 2011. Food animals and antimicrobials: impacts on human health. Clin Microbiol Rev 24: 718–733.

11. Robinson TP, Bu DP, Carrique-Mas J, Fèvre EM, Gilbert M, Grace D, Hay SI, Jiwakanon J, Kakkar M, Kariuki S, Laxminarayan R, Lubroth J, Magnusson U, Thi Ngoc P, Van Boeckel TP, Woolhouse ME. 2016. Antibiotic resistance is the quintessential One Health issue. Trans R Soc Trop Med Hyg 110: 377–380.

12. Alvarez-Uria G, Gandra S, Laxminarayan R. 2016. Poverty and prevalence of antimicrobial resistance in invasive isolates. Int J Infect Dis 52 SRC-:59–61.

13. Furtula V, Farrell E, Diarrassouba F, Rempel H, Pritchard J, Diarra M. 2010. Veterinary pharmaceuticals and antibiotic resistance of *Escherichia coli* isolates in poultry litter from commercial farms and controlled feeding trials. Poult Sci 89: 180–188.

14. Vasco K, Graham JP, Trueba G. 2016. Detection of zoonotic enteropathogens in children and domestic animals in a semi-rural community in Ecuador. Appl Environ Microbiol 82: 4218–4224.

15. Graham DW, Knapp CW, Christensen BT, McCluskey S, Dolfing J. 2016. Appearance of β-lactam resistance genes in agricultural soils and clinical isolates over the 20th Century. Sci Rep 6: 21550.

16. Zhang H, Zhai Z, Li Q, Liu L, Guo S, Yang L, Ye C, Chang W, Zhai J. 2016. Characterization of extended-spectrum β-lactamase-producing *Escherichia coli* isolates from pigs and farm workers. J Food Prot 79:1630–1634.

17. Lowenstein C, Waters WF, Roess A, Leibler JH, Graham JP. 2016. Animal husbandry practices and perceptions of zoonotic infectious disease risks among livestock keepers in a rural parish of Quito, Ecuador. Am J Trop Med Hyg 95: 1450–1458.

18. Roess AA, Winch PJ, Ali NA, Akhter A, Afroz D, El Arifeen S, Darmstadt GL, Baqui AH. 2013. Animal husbandry practices in rural Bangladesh: Potential risk factors for antimicrobial drug resistance and emerging diseases. Am J Trop Med Hyg 89: 965–970.

19. Pehrsson EC, Tsukayama P, Patel S, Mejía-Bautista M, Sosa-Soto G, Navarrete KM, Calderon M, Cabrera L, Hoyos-Arango W, Bertoli MT, Berg DE, Gilman RH, Dantas G. 2016. Interconnected microbiomes and resistomes in low-income human habitats. Nature 533:212–216.

20. Blyton MD, Pi H, Vangchhia B, Abraham S, Trott DJ, Johnson JR, Gordon DM. 2015. Genetic structure and antimicrobial resistance of *Escherichia coli* and cryptic clades in birds with diverse human associations. Appl Environ Microbiol 81: 5123–5133.

21. Landers TF, Cohen B, Wittum TE, Larson EL. 2012. A review of antibiotic use in food animals: Perspective, policy, and potential. Public Health Rep 127: 4–22.

22. Andersson DI. 2003. Persistence of antibiotic resistant bacteria. Curr Opin Microbiol 6: 452–456.

23. Magiorakos AP, Srinivasan A, Carey R, Carmeli Y, Falagas M, Giske C, Harbarth S, Hindler J, Kahlmeter G, Liljequist B. 2012. Olsson-Multidrug-resistant, extensively drug-resistant and pandrug-resistant bacteria: an international expert proposal for interim standard definitions for acquired resistance. Clin Microbiol Infect 18: 268–281.

24. Moeller AH, Suzuki TA, Phifer-Rixey M, Nachman MW. 2018. Transmission modes of the mammalian gut microbiota. Science 362: 453–457.

25. Lautenbach E, Bilker WB, Tolomeo P, Maslow JN. 2008. Impact of diversity of colonizing strains on strategies for sampling *Escherichia coli* from fecal specimens. J Clin Microbiol 46: 3094–3096.

26. Patel J, Cockerill F, Alder J, Bradford P, Eliopoulos G, Hardy D. 2014. Performance standards for antimicrobial susceptibility testing; twenty-fourth informational supplement. CLSI Stand Antimicrob susceptibility Test 34:1–226.

27. Miller JH. A short course in bacterial genetics: a laboratory manual and handbook for *Escherichia coli* and related bacteria. Trends in Biochemical Sciences-Library Compendium. 1993;18:193. 28.

28. Jacoby GA, Chow N, Waites KB. 2003. Prevalence of plasmid-mediated quinolone resistance. Antimicrob Agents Chemother 47: 559–562.

29. Carattoli A. 2013. Plasmids and the spread of resistance. Int J Med Microbiol 303: 298–304.

30. Bolger Anthony M., Marc L, Bjoern U. 2014. Trimmomatic: A flexible trimmer for Illumina Sequence Data. Bioinformatics.

31. Bankevich A, Nurk S, Antipov D, Gurevich AA, Dvorkin M, Kulikov AS, Lesin VM, Nikolenko SI, Pham S, Prjibelski AD, Pyshkin A V, Sirotkin A V, Vyahhi N, Tesler G, Alekseyev MA, Pevzner PA. 2012. SPAdes: a new genome assembly algorithm and its applications to single-cell sequencing. J Comput Biol 19: 455–477.

32. Koren S, Walenz BP, Berlin K, Miller JR, Bergman NH, Phillippy AM. 2017. Canu: scalable and accurate long-read assembly via adaptive k-mer weighting and repeat separation. Genome Res 27:722–736.

33. Seemann T. 2014. Prokka: rapid prokaryotic genome annotation. Bioinformatics 30: 2068–2069.

34. Center for Genomic Epidemiology. ResFinder 2.1 / PlasFinder 1.3 / pMLST 1.4 / MLST 1.8.

35. Kumar S, Stecher G, Tamura K. 2016. MEGA molecular evolutionary genetics analysis version 7. 0 bigger datasets Mol Biol Evol 33: 1870–1874.

36. Darling AE, Tritt A, Eisen JA, Facciotti MT. 2011. Mauve assembly metrics. Bioinformatics 27: 2756–2757.

37. Edwards DJ, Holt KE. 2013. Beginner’s guide to comparative bacterial genome analysis using next-generation sequence data. Microb informatics Exp 2 3 SRC-B.

38. Nascimento M, Sousa A, Ramirez M, Francisco AP, Vaz C. 2016. Carriço JA, PHYLOViZ 2. Provid scalable data Integr Vis Mult phylogenetic inference methods Bioinforma 33: 128–129.

39. Siguier P, Lestrade L, Mahillon J, Chandler M. 2006. Pérochon J, ISfinder: the reference centre for bacterial insertion sequences. Nucleic Acids Res 34: D32–D36.

40. Gouliouris T, Raven KE, Ludden C, Blane B, Corander J, Horner CS, Hernandez-Garcia J, Wood P, Hadjirin NF, Radakovic M. 2018. Genomic surveillance of *Enterococcus faecium* reveals limited sharing of strains and resistance genes between livestock and humans in the united kingdom. MBio 9: e01780–18.

41. Johnson JR, Clabots C, Kuskowski MA. 2008. Multiple-host sharing, long-term persistence, and virulence of Escherichia coli clones from human and animal household members. J Clin Microbiol 46: 4078–4082.

42. Johnson JR, Miller S, Johnston B, Clabots C, DebRoy C. 2009. Sharing of Escherichia coli sequence type ST131 and other multidrug-resistant and urovirulent E. coli strains among dogs cats within a Househ J Clin Microbiol 47: 3721–3725.

43. Johnson JR, Clabots C. 2006. Sharing of virulent Escherichia coli clones among household members of a woman with acute cystitis. Clin Infect Dis 43: e101–e108.

44. Tartof SY, Solberg OD, Manges AR, Riley LW. 2005. Analysis of a uropathogenic Escherichia coli clonal group by multilocus sequence typing. J Clin Microbiol 43: 5860–5864.

45. Mushtaq S, Irfan S, Sarma J, Doumith M, Pike R, Pitout J, Livermore D, Woodford N. 2011. Phylogenetic diversity of *Escherichia coli* strains producing NDM-type carbapenemases. J Antimicrob Chemother 66 :2002–2005.

46. Lau SH, Reddy S, Cheesbrough J, Bolton FJ, Willshaw G, Cheasty T, Fox AJ, Upton M. 2008. Major uropathogenic *Escherichia coli* strain isolated in the northwest of England identified by multilocus sequence typing. J Clin Microbiol 46: 1076–1080.

47. de Been M, Lanza VF, de Toro M, Scharringa J, Dohmen W, Du Y, Hu J, Lei Y, Li N, Tooming-Klunderud A. 2014. Dissemination of cephalosporin resistance genes between *Escherichia coli* strains from farm animals and humans by specific plasmid lineages. PLoS Genet 10: e1004776.

48. Roetzer A, Diel R, Kohl TA, Blom J, Wirth T, Jaenicke S, Schuback S, Gerdes S. 2013. Rückert C, Nübel U, Rüsch-Whole genome sequencing versus traditional genotyping for investigation of a *Mycobacterium tuberculosis* outbreak: a longitudinal molecular epidemiological study. PLoS Med e1001387

49. Guo J, Li J, Chen H, Bond PL, Yuan Z. 2017. Metagenomic analysis reveals wastewater treatment plants as hotspots of antibiotic resistance genes and mobile genetic elements. Water Res 123: 468–478.

50. Woolhouse M, Ward M, van Bunnik B, Farrar J, Society B. 2014. Antimicrobial resistance in humans, livestock and the wider environment. Philos Trans R Soc Lond B Biol Sci 370: 20140083

51. Stokes HW, Gillings MR. 2011. Gene flow, mobile genetic elements and the recruitment of antibiotic resistance genes into Gram-negative pathogens. FEMS Microbiol Rev 35:790–819.

52. Fang L-X, Li X-P, Deng G-H, Li S-M, Yang R-S, Wu Z-W, Liao X-P, Sun J, Liu Y-H. 2018. High genetic plasticity in multidrug-resistant sequence type 3-IncHI2 plasmids revealed by sequence comparison and phylogenetic analysis. Antimicrob Agents Chemother 62: e02068–17.

